# Identifying potential risk genes for clear cell renal cell carcinoma with deep reinforcement learning

**DOI:** 10.1101/2024.06.19.599667

**Authors:** Dazhi Lu, Yan Zheng, Jianye Hao, Xi Zeng, Lu Han, Zhigang Li, Shaoqing Jiao, Jianzhong Ai, Jiajie Peng

## Abstract

Clear cell renal cell carcinoma (ccRCC) is the most prevalent type of renal cell carcinoma. However, our understanding of ccRCC risk genes remains limited. This gap in knowledge poses significant challenges to the effective diagnosis and treatment of ccRCC. To address this problem, we propose a deep reinforcement learning-based computational approach named RL-GenRisk to identify ccRCC risk genes. Distinct from traditional supervised models, RL-GenRisk frames the identification of ccRCC risk genes as a Markov decision process, combining the graph convolutional network and Deep Q-Network for risk gene identification. Moreover, a well-designed data-driven reward is proposed for mitigating the limitation of scant known risk genes. The evaluation demonstrates that RL-GenRisk outperforms existing methods in ccRCC risk gene identification. Additionally, RL-GenRisk identifies ten novel ccRCC risk genes. We successfully validated epidermal growth factor receptor (EGFR), corroborated through independent datasets and biological experimentation. This approach may also be used for other diseases in the future.

## 1 Introduction

Renal cell carcinoma (RCC), one of the most common cancers worldwide, is a type of kidney cancer that initiates in the lining of the proximal convoluted tubule [1, 2]. Clear cell renal cell carcinoma (ccRCC) constitutes 80% of all RCC cases and is particularly aggressive due to its high immune infiltration [3–5]. In addition, over 30% of ccRCC patients suffer from metastasis, which is a significant factor leading to death in ccRCC patients [6–8]. Although several drugs have been utilized for the treatment of ccRCC [9, 10], the efficacy is still limited due to the heterogeneity of ccRCC [11]. Therefore, it is necessary to understand the pathogenesis and identify risk genes of ccRCC, which may be beneficial for early diagnosis and treatment of ccRCC [12–14].

Cancer is a complex genetic disorder. Its occurrence and progression are associated with the accumulation of driver genetic mutations that provide a selective growth advantage to cells [15, 16]. Consequently, one class of methods to identify cancer risk genes is based on mutation data. In the past years, several cancer sequencing projects have generated mutation data from thousands of cancer patients, enhancing the identification of cancer risk genes [15, 17]. Traditional statistical approaches focus on genes with a higher mutation frequency in the patient cohort than the control cohort [18]. Youn et al. [19] employed the functional impact of mutations on proteins, variations in background mutation rates among tumors, and the redundancy of the genetic code in tumor genome sequencing data to identify key genes in non-small cell lung cancer. Methods like MuSiC [20], OncodriveCLUST [21], and MutSigCV [22] identified cancer risk genes by comparing the observed gene mutations with the predefined background mutation frequency. By now, frequency-based methods have identified many cancer risk genes and enhanced cancer diagnosis and therapy [23]. However, the genetic foundations of cancer are highly diverse. Except for genes mutated across a large number of patients, some key genes in tumor initiation and progression are observed to be mutated in only a few patients [24]. For example, PIK3CA, which has been validated to be a ccRCC risk gene by previous studies [25–27], is mutated in no more than 5% of ccRCC patients [27]. It is difficult for purely frequency-based approaches to identify these genes with low mutation frequency but high risk.

To address the drawback of frequency-based methods, the interactions among proteins are introduced for cancer risk gene identification, since genes involved in the same signaling and regulatory pathways as well as protein complexes may interact to exert their effects together. Muffinn [28] identified cancer risk genes through network propagation, taking into account mutations not only in individual genes but also in their neighbors within the protein-protein interaction (PPI) network. DiSCaGe [29] calculated a gene mutation score using an asymmetric spreading strength based on the type of mutations and the PPI network, then produced a ranking of prioritized cancer risk genes. HotNet2 [29] used an insulated heat diffusion process to identify cancer risk genes by propagating heat through the PPI network. nCOP [30] employed a heuristic search method to select connected subnetworks from the PPI network based on the mutation data of cancer patients, and then ranked cancer risk genes based on the frequencies of genes appearing in these subnetworks. Both aforementioned methods are unsupervised, which may suffer from the highly diverse genetic foundations of cancer or the noise in the PPI network [31]. Recently, several supervised methods have emerged as potentially valuable tools for predicting cancer risk genes [23, 32–35]. For example, Agajanian Steve et al. [33] trained a random forest classifier to identify cancer risk genes based on the known cancer-driver mutations. DeepDriver [34] used gene mutation types as features, constructed a K-nearest neighbor graph based on the Pearson correlation coefficient, and trained a convolutional neural network to identify cancer risk genes. Nevertheless, different from unsupervised methods [28–30, 36], supervised methods require a substantial amount of known high-confidence risk genes as labeled data for model training [23]. Unfortunately, the number of known high-confidence ccRCC risk genes is currently limited [37, 38]. For example, there are only 27 ccRCC risk genes in the IntOGen database [39]. Owing to the reliance on labels, supervised methods may not be suitable for identifying ccRCC risk genes.

To overcome the limitations of existing methods, we propose a deep reinforcement learning-based approach for ccRCC risk gene identification, named RL-GenRisk (Rein-forcement Learning-based GENe RISK). The reinforcement learning-based model leverages environmental interactions for optimization [40], tackling the challenge of scant known risk genes. Specifically, RL-GenRisk models the PPI network as the environment and utilizes a graph convolutional network [41] to learn state representations. It also incorporates the Deep Q-Network (DQN) [42] to combine reinforcement learning with deep neural networks for ccRCC risk gene identification. Moreover, a data-driven reward is designed to facilitate a straightforward method for identifying ccRCC risk genes. By focusing on a sampled subgraph with node features, the data-driven reward effectively leverages information from both the PPI network and gene mutation data. This not only ensures the accurate identification of genes with high mutation frequencies but also enables RL-GenRisk to identify potential risk genes with low mutation frequencies that functionally interact with genes having high mutation frequencies. Extensive experiments demonstrate that RL-GenRisk outperforms the existing methods in the identification of ccRCC risk genes. Furthermore, several novel risk genes are revealed and validated in independent datasets. Specifically, we validated a high-confidence gene EGFR through statistical, *in vitro*, and *in vivo* experiments. Statistical analyses show significant upregulation of EGFR at both bulk and single-cell levels among ccRCC patients, with a significant association between over-expression of the protein encoded by EGFR and poor survival in ccRCC patients. The *in vitro* experimental results show that decreased EGFR expression promotes ccRCC cell apoptosis as well as suppresses colony formation and migration, and the use of the EGFR inhibitor erlotinib effectively augments apoptosis and inhibits migration. Moreover, the *in vivo* experimental results show that both the erlotinib and EGFR downregulation can significantly repress the growth of ccRCC tumors in mice.

## 2 Results

### 2.1 RL-GenRisk framework

We propose a deep reinforcement learning-based approach to identify ccRCC risk genes (Fig 1), named RL-GenRisk. Fundamentally different from existing supervised deep learning-based methods, RL-GenRisk incorporates the reinforcement learning paradigm. Specifically, RL-GenRisk frames the ccRCC risk gene identification as a sequence decision-making process, formulated as a Markov decision process [43]. This enables RL-GenRisk to seamlessly integrate reinforcement learning algorithms, thereby effectively addressing the inherent challenge of scant known risk genes. RL-GenRisk takes PPI network and gene mutation data as input (Fig 1. A). The PPI network is represented as an undirected graph, with nodes representing genes and edges representing interactions between genes. Gene mutation data includes details on the presence of mutations in ccRCC patients for each gene. RL-GenRisk treats the PPI network with gene mutation information as the environment. The state includes a sampled subgraph with node features. The action is selecting a node directly connected to the sampled subgraph and adding it to this sampled subgraph. Thus, the risk gene identification is framed as a Markov decision process of node selection within the PPI network.

**Fig. 1.**
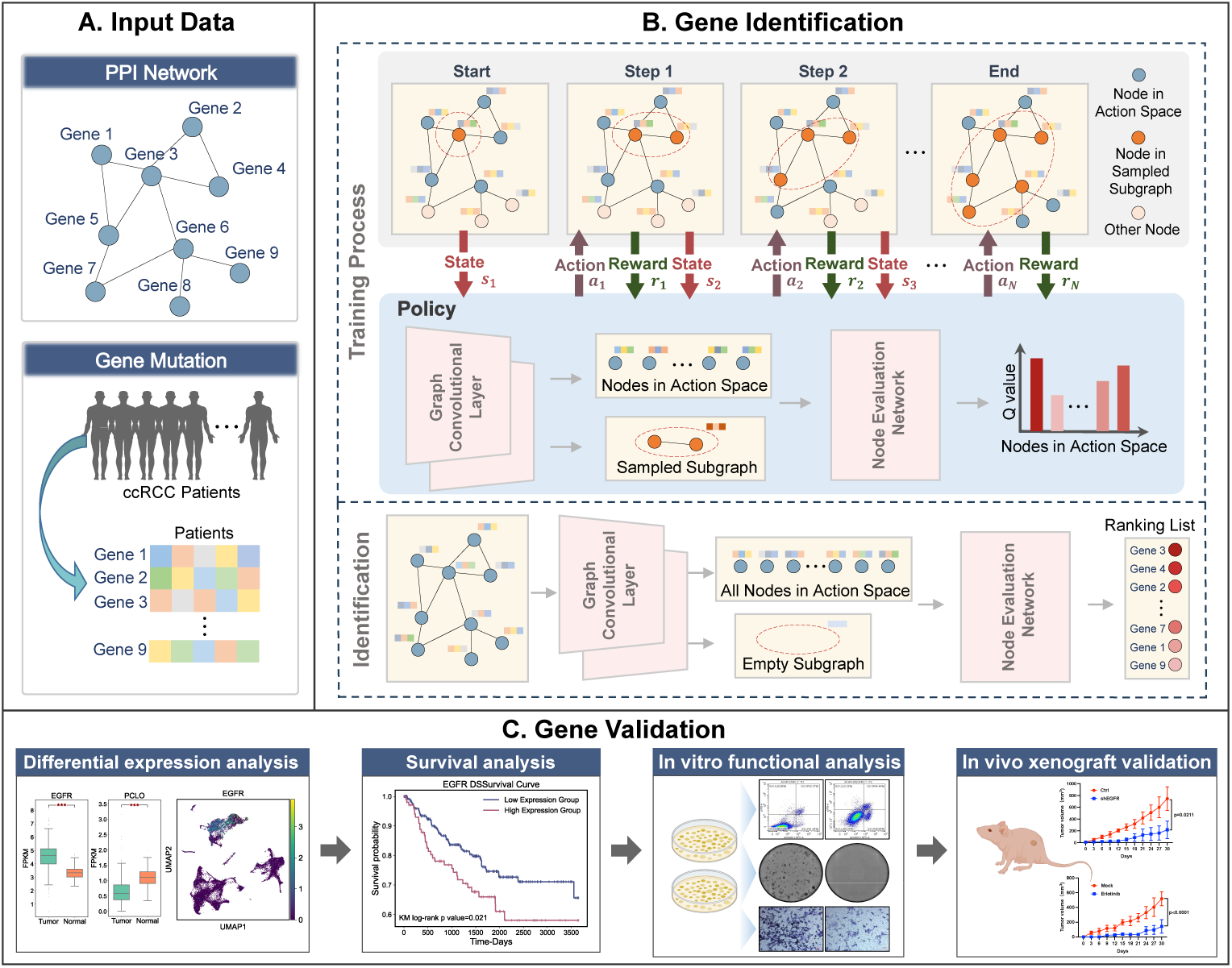
Overview of workflow. **A**, Input data of RL-GenRisk, including a Protein-Protein Inter-action (PPI) network and gene mutation data of ccRCC patients from The Cancer Genome Atlas (TCGA). **B**, Overview of the training and identification process. During the training process, RL-GenRisk starts by selecting a node randomly and appending this node to a sampled subgraph (indicated by the red dashed circle). Nodes colored orange are included in the sampled subgraph. Nodes interacting with the sampled subgraph are colored blue, indicating their inclusion in the action space. The policy comprises two graph convolutional layers and a node evaluation network. It takes the current state as input and selects an action based on *Q* values of nodes in the action space. Subsequently, the state is updated, and the reward is obtained. This process repeats until the sampled subgraph reaches its maximum size. Following training, RL-GenRisk computes *Q* values for all genes based on an empty subgraph, resulting in a ranking list of risk genes ordered by *Q* values. **C**, Experiment validation on external data for identified ccRCC risk genes.

Central to RL-GenRisk is a policy that is represented by a neural network and interacts with the environment. The policy takes the current state as input and predicts *Q* values, representing the probability distribution of all possible actions. The policy of RL-GenRisk consists of two main components: a Graph Convolutional Network (GCN) for learning state representation and a node evaluation network for computing action probability (Fig 1. B top). To enhance state representation, RL-GenRisk employs the GCN [41], an inductive graph representation learning method, to capture node representations in the PPI network. Node initial features are derived from the PPI network’s topology information and ccRCC patients’ mutation information. The policy of RL-GenRisk is trained to select the optimal actions by maximizing the reward. In this study, we designed a data-driven reward that focused on the sampled subgraph, considering both information from the PPI network and gene mutation data. The DQN algorithm is employed to update the policy’s parameters. Throughout the training, we employed the *ɛ*-greedy strategy to choose actions based on *Q* values, thus boosting RL-GenRisk’s exploratory potential. In the identification phase (Fig 1. B bottom), RL-GenRisk starts with an empty subgraph, incorporates all nodes into the action space, and utilizes the trained policy to calculate *Q* values for each node. Subsequently, ccRCC risk genes are ranked by *Q* values, with higher values indicating greater risk. Further details about the RL-GenRisk can be found in the “Methods” section.

### 2.2 RL-GenRisk shows superior performance for ccRCC risk gene prioritization over existing methods

To assess the performance of RL-GenRisk, we utilized RL-GenRisk and five other existing methods to identify ccRCC genes with gene mutation data of ccRCC patients from The Cancer Genome Atlas (TCGA) [44] and five different PPI networks, including HPRD [45], STRING-db [46], Multinet [47], IRefIndex [48] and HumanNet [49]. Known ccRCC risk genes were retrieved from the IntOGen [39] cancer-specific database. We employed the Discounted Cumulative Gain (DCG) as the evaluation metric, consistent with prior research [29, 50]. Specifically, the DCG scores for the top 100 genes identified by each method were calculated. Following this, we calculated the area under the DCG curve (DCG-AUC) to further assess these methods’ predictive accuracy. Evaluation results indicated that RL-GenRisk outperformed the five other established methods in identifying ccRCC risk genes, achieving the topmost DCG score of 5.29 (as shown in Supplementary Figure 1. A) and DCG-AUC score of 4.49 (as shown in Supplementary Figure 1. B). As anticipated, MutsigCV had the lowest performance due to its reliance solely on mutation data, lacking integration with biological network insights. Additionally, RL-GenRisk demonstrated more stable predictive capabilities across different PPI networks compared to the other four methods that also utilize PPI networks (Fig 2. B and Fig 2. C). RL-GenRisk with HPRD achieved the best performance among all methods across different PPI networks (Fig 2. A and Supplementary Figure 2). Muffinn excelled on HumanNet but showed inferior performance on other PPI networks, highlighting its instability across different PPI networks. HotNet2’s DCG-AUC scores across various PPI networks varied greatly, suggesting significant inconsistencies in its ranking predictions of known ccRCC genes across different PPI networks. Conversely, while nCOP and DiSCaGe showed stable performance across PPI networks, their overall efficacy was found to be inferior compared to RL-GenRisk. The subsequent analyses were grounded on the high-confidence risk genes (HRGs) identified by the best-performing method, RL-GenRisk with the HPRD network.

**Fig. 2.**
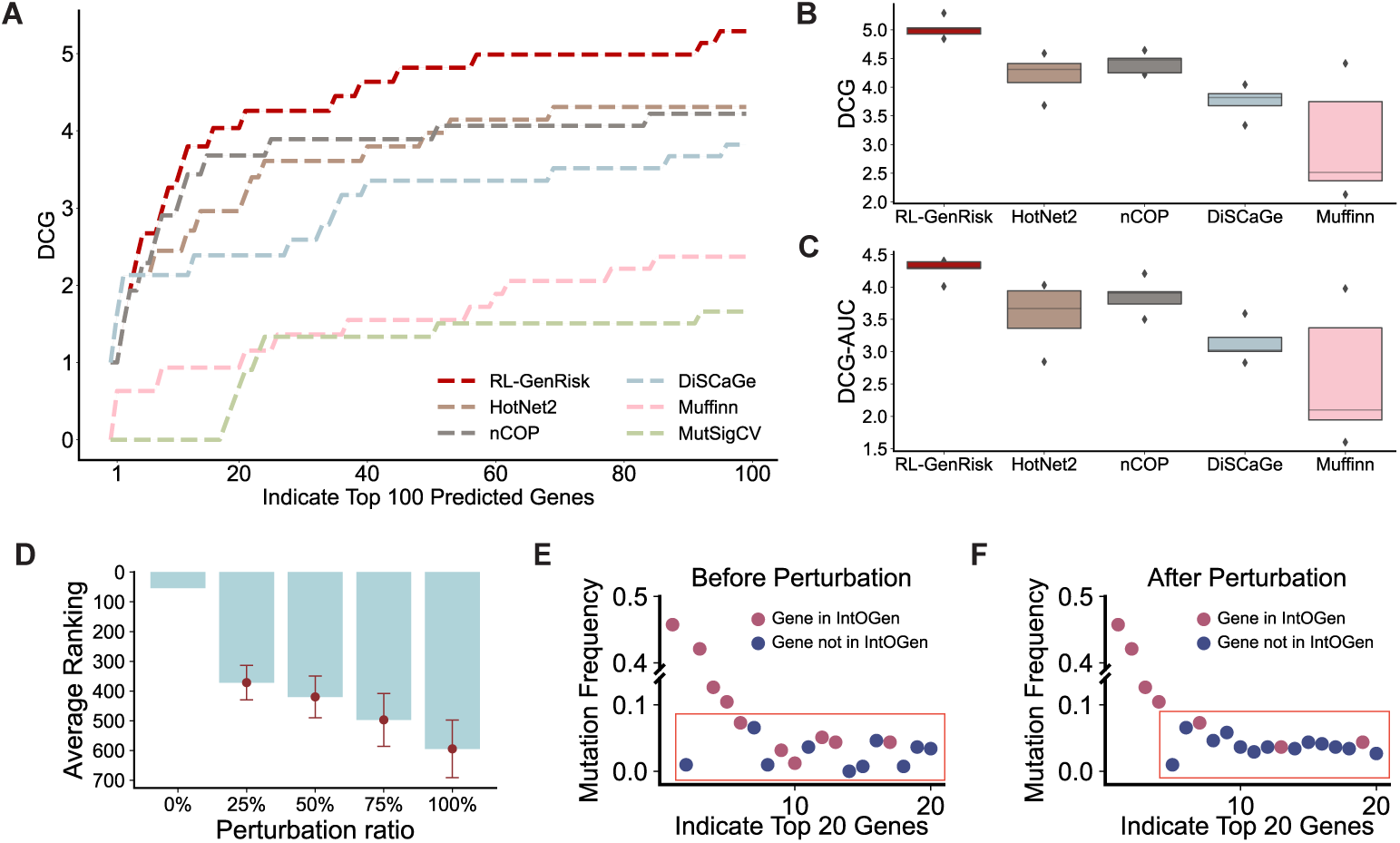
Performance comparison and perturbation analysis. **A**. Discounted Cumulative Gain (DCG) curves of the top 100 identified genes with RL-GenRisk and other compared methods on the HPRD network. **B, C**. Boxplot of DCG scores and area under DCG curve (DCG-AUC) on different PPI networks across different methods. **D**. Average ranking of known ccRCC genes with low mutation frequency (less than 5%) predicted by RL-GenRisk after systematic perturbation of the PPI network. **E, F.** Mutation frequency of the top 20 genes identified by RL-GenRisk (E. Before perturbation; F. After perturbation.). Each dot represents a gene. Dot colored red indicates that the gene is included in the IntOGen database, and colored blue indicates that the gene is not included in the IntOGen database. And the mutation frequency of the genes enclosed by the orange box is less than 10%.

### 2.3 Biological network facilitates the identification of low-frequency mutated ccRCC risk genes

Although gene mutation frequency is crucial for assessing cancer associations, not all cancer-related genes have high mutation frequencies [51]. Therefore, relying solely on mutation data might overlook low mutation frequency, high-risk cancer genes. To evaluate the capability of RL-GenRisk in identifying ccRCC risk genes with low mutation frequency but high risk, we utilized the “maftools” package [52] to analyze the mutation frequencies of the top 20 HRGs (Supplementary Table 1) identified by RL-GenRisk in ccRCC patients from TCGA (Supplementary Figure 3). Generally, 315 (76.64%) of the 411 ccRCC patients from TCGA exhibited somatic mutations in the top 20 HRGs. Missense mutations were predominant, followed by frameshift mutations, among the identified somatic mutation types. Notably, the mutation frequencies of four genes, including VHL, PBRM1, SETD2, and BAP, exceeded 10%. These genes are already recognized as ccRCC risk genes and included in the IntOGen database. Genes with high mutation frequencies are more likely to be detected by methods that solely rely on mutation data. However, it’s important to note that not all cancer genes have high mutation frequencies [51]. Among the top 20 HRGs, five recognized ccRCC risk genes in the IntOGen database had mutation frequencies below 5%. For example, only 5 of 411 ccRCC patients carry mutations in PIK3CA, a known ccRCC risk gene in the IntOGen database, indicating variable mutation frequencies among cancer genes. To assess the contribution of the biological network in RL-GenRisk for identifying low mutation frequency, high-risk ccRCC genes, we progressively increased the percentage of edges randomly swapped between node pairs from 25% up to 50%, 75%, and 100%. Perturbing the PPI network disrupts its internal information, with more extensive perturbation causing greater information loss. As the perturbation ratio increased, the rankings of known ccRCC genes that initially had high ranks and mutation fre-quencies below 5% gradually dropped (as shown in Fig 2. D). We then compared the mutation frequencies of the top 20 HRGs identified by RL-GenRisk before and after PPI network perturbation and created dot plots for visualization (Fig 2. E and Fig 2. F). We found that after network perturbation, three known ccRCC risk genes, TP53, PIK3CA, and SPEN, previously identified in the top 20 with mutation frequencies below 3%, dropped out of the top 20. This indicates that incorporating PPI net-work knowledge improves the detection of low mutation frequency ccRCC genes with significant risk implications.

### 2.4 Biological function analysis of high-confidence risk genes

To delve into the biological function of HRGs, we conducted pathway enrichment analysis using the WikiPathways database [53], a comprehensive resource for pathway-based data analysis. Enrichment analysis was performed using g:Profiler [54]. Fig 3. A illustrated the top 10 significantly enriched pathways (FDR p-value *<*= 1.21e-4). The results demonstrated significant enrichment in various cancer-related pathways, with “Clear cell renal cell carcinoma pathways” being the most prominently enriched (FDR p-value = 1.48e-8). The Human Phenotype Ontology (HPO) database [55], which pro-vides a standardized vocabulary of phenotypic abnormalities associated with human diseases, highlighted “Renal neoplasm” and “Renal cell carcinoma” as the most sig-nificantly enriched phenotypes (FDR p-value *<* 4.05e-5, Supplementary Figure 4). Furthermore, Gene Ontology (GO) enrichment analysis indicated that HRGs were notably enriched in various cancer-related biological processes, including “cell adhe-sion” (FDR p-value = 2.69e-4), “regulation of cell population proliferation” (FDR p-value = 3.64e-4), “cell population proliferation” (FDR p-value = 6.85e-4), and “cell migration” (FDR p-value = 2.39e-3). These observations showed a comprehensive molecular landscape linked to HRGs within the scope of ccRCC, emphasizing the critical roles of these genes in ccRCC-related mechanisms.

**Fig. 3.**
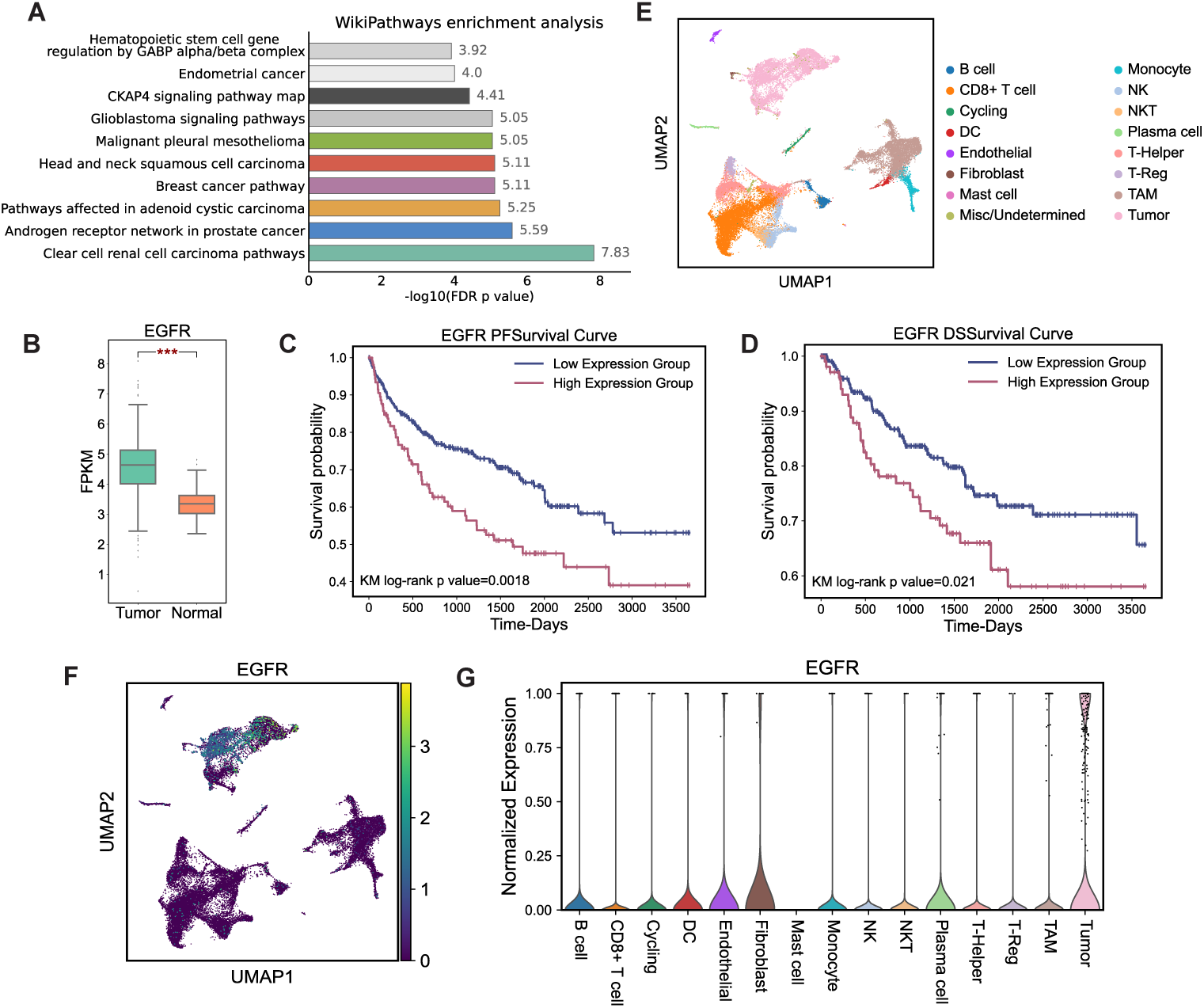
Independent datasets analysis. **A**. Top 10 enriched pathways of the Wikipathways based on the top 20 high-confidence genes (HRGs). **B**. The expression levels of EGFR in tumor and normal tissues of ccRCC patients from TCGA, quantified using Fragments Per Kilobase Million (FPKM), “***” indicates p-value *<* 0.001. **C, D**. Survival curves of EGFR. ccRCC patients from TCGA were divided into two groups based on the top 25% and bottom 25% of expression levels of protein encoded by EGFR, using Progression-Free Survival (PFS) and Disease-Specific Survival (DSS) respectively. The red curve corresponds to the high expression group, while the blue curve corresponds to the low expression group. **E**. UMAP plot of clustering of single-cell RNA-seq data from kidneys of ccRCC patients. DC, dendritic cell; NK, natural killer cell; NKT, natural killer T cell; TAM, tumor-associated macrophage; T-Reg, regulatory T cell. **F**. Gene expression of EGFR in different cells of ccRCC patients, with colors corresponding to the expression values. **G**. Violin plot displaying the normalized gene expression of EGFR in different cells of ccRCC patients in the single-cell RNA-seq data.

### 2.5 Differential expression analysis revealed significant differential gene expression of EGFR at both bulk and single-cell levels

Among the top 20 HRGs identified by RL-GenRisk, 10 HRGs are newly identi-fied ccRCC risk genes, which are not included in the IntOGen database. We then performed differential expression analysis on these 10 newly identified HRGs using RNA-seq data from 607 ccRCC patients in TCGA. Notably, EGFR and PCLO showed significant differential expression between tumor tissues and normal tissues among these 10 newly identified HRGs (p-value *<* 1.98e-21, Wilcoxon signed-rank test, Fig 3. B and Supplementary Figure 5). The differential expression results for these 10 newly identified HRGs are shown in Supplementary Figure 5. EGFR was notably upregulated in ccRCC tumor tissues compared to normal tissues. EGFR is activated as a homodimer or heterodimer, thereby regulating multiple signaling pathways, including the RAS/RAF/MAPK, AKT, and JAK/STAT pathways, which play essential roles in cell migration, proliferation, and survival [56–58]. These pathways are intricately involved in driving cell proliferation and conferring resistance to apoptosis [58]. The overexpression of EGFR leads to an excess of receptors on the cell surface, fostering uncontrolled cell growth and division. This dysregulation can drive the transformation of normal cells into tumor cells, creating a favorable environment for sustained tumor cell survival [59]. Additionally, the protein encoded by PCLO is a component of the presynaptic cytoskeletal matrix, which is involved in establishing active synaptic zones and in synaptic vesicle trafficking [60]. Recent research has identified the expression level of PCLO as a prognostic biomarker for esophageal squamous cell carcinoma [61], and a notable high mutation frequency (47.9%) of PCLO has been observed in large Central European cohorts with gastric cancer [62]. In our study, the observed elevated expression of PCLO in normal tissues relative to tumor tissues suggests the potential of PCLO to act as a tumor suppressor or modulator in the progression of cancer.

To further investigate the expression patterns of EGFR and PCLO in tumor cells and normal cells of ccRCC patients, we performed an analysis utilizing single-cell RNA-seq data from seven ccRCC patients reported in a recent work [63] (Supplementary Table 2). Uniform manifold approximation and projection (UMAP) was used to visualize the distribution of 31,856 cells from kidneys in these patients in a two-dimensional plane (Fig 3. E). We used UMAP to visualize the expression of the top 20 HRGs in different cells (Supplementary Figure 6) and plotted the distribution of expression of these genes in different cell types (Supplementary Figure 7). Subsequently, we assessed the expression levels of EGFR and PCLO across different cell types within these patients (Fig 3. F and Supplementary Figure 6). Our investigation showed a significant difference in EGFR expression between tumor cells and other cells, (p-value *<*= 8.41e-5, Wilcoxon signed-rank test, Supplementary Figure 8), with displaying higher expression levels in tumor cells (Fig 3. G). On the other hand, while PCLO exhibited significant differential expression in bulk data between tumor and normal tissues, no significant differential expression was observed at the single-cell level (Supplementary Figure 5 and Supplementary Figure 8). Therefore, the significant upregulation of EGFR at both bulk and single-cell levels in ccRCC patients suggested its potential as a biomarker for ccRCC.

### 2.6 Expression level of protein encoded by the EGFR is significantly correlated with the prognosis of ccRCC patients

To reveal whether EGFR affects the prognosis of ccRCC patients, the survival analysis is conducted utilizing clinical data and expression of EGFR-encoded protein obtained from the TCGA dataset. We obtained clinical data and reverse-phase protein array (RPPA) data of ccRCC patients from TCGA. Progression-Free Survival (PFS) [64] and Disease-Specific Survival (DSS) [65] were utilized to assess the relationship between the expression levels of protein encoded by EGFR and the survival of ccRCC patients. The ccRCC patients from TCGA were categorized into quartiles based on the expression levels of protein encoded by EGFR. We analyzed the ten-year survival rate following cancer diagnosis and plotted Kaplan-Meier survival curves to illustrate the impact of EGFR encoded protein expression on the prognosis of patients (Fig 3. C and Fig 3. D). The results of the survival analysis revealed a significant association between the expression levels of protein encoded by EGFR and the survival time of ccRCC patients, with higher EGFR encoded protein expression being correlated with poorer survival outcomes (KM log-rank p-value = 0.0018, PFS; KM log-rank p-value = 0.021, DSS). Our observations revealed that the overexpression of protein encoded by EGFR may play a critical role in cancer progression and could potentially serve as a prognostic biomarker for ccRCC patients.

### 2.7 EGFR effectively promotes ccRCC progression *in vitro* and *in vivo*

To verify the effect of EGFR expression on ccRCC progression, the stable cells harboring EGFR knockdown were obtained using ACHN (Fig 4. A and Fig 4. B) and 786-O cells (Fig 4. G and Fig 4. H). The protein and mRNA expression of EGFR was quantitated by western blotting and qPCR, respectively, and short hairpin RNA (shRNA)-2/3 showed promising knockdown efficacies for EGFR silencing (Fig 4. A, B, G, and H). The downregulation of EGFR significantly inhibits the cell viability (CCK8 assay, Fig 4. C-ACHN, Fig 4. I-786-O) and migration (transwell assay, Fig 4. E-ACHN, Fig 4. K-786-O). Furthermore, decreased EGFR expression markedly promotes the ccRCC cell apoptosis (flow cytometry assay, Fig 4. D-ACHN, Fig 4. J-786-O) and represses cell colony formation *in vitro* (colony assay, Fig 4. F-ACHN, Fig 4. L-786-O). Also, the EGFR overexpression was detected using qPCR (Fig 4.M) and western blot-ting (Fig 4. N), and it promotes the migration of 786-O cells significantly (Fig 4. O). Moreover, the EGFR inhibitor, erlotinib, was further used to inhibit its activity, and the data indicated that erlotinib can effectively inhibit ccRCC cell migration (Fig 5. A middle panel) and growth (Fig 5. A lower panel) as well as promotes the apoptosis (Fig 5. A upper panel) *in vitro*. *In vivo*, both erlotinib and EGFR downregulation can markedly repress the tumor growth (Fig 5. B and Fig 5. C). Taken together, these findings suggested that EGFR can significantly promote ccRCC cell progression as a risk factor.

**Fig. 4.**
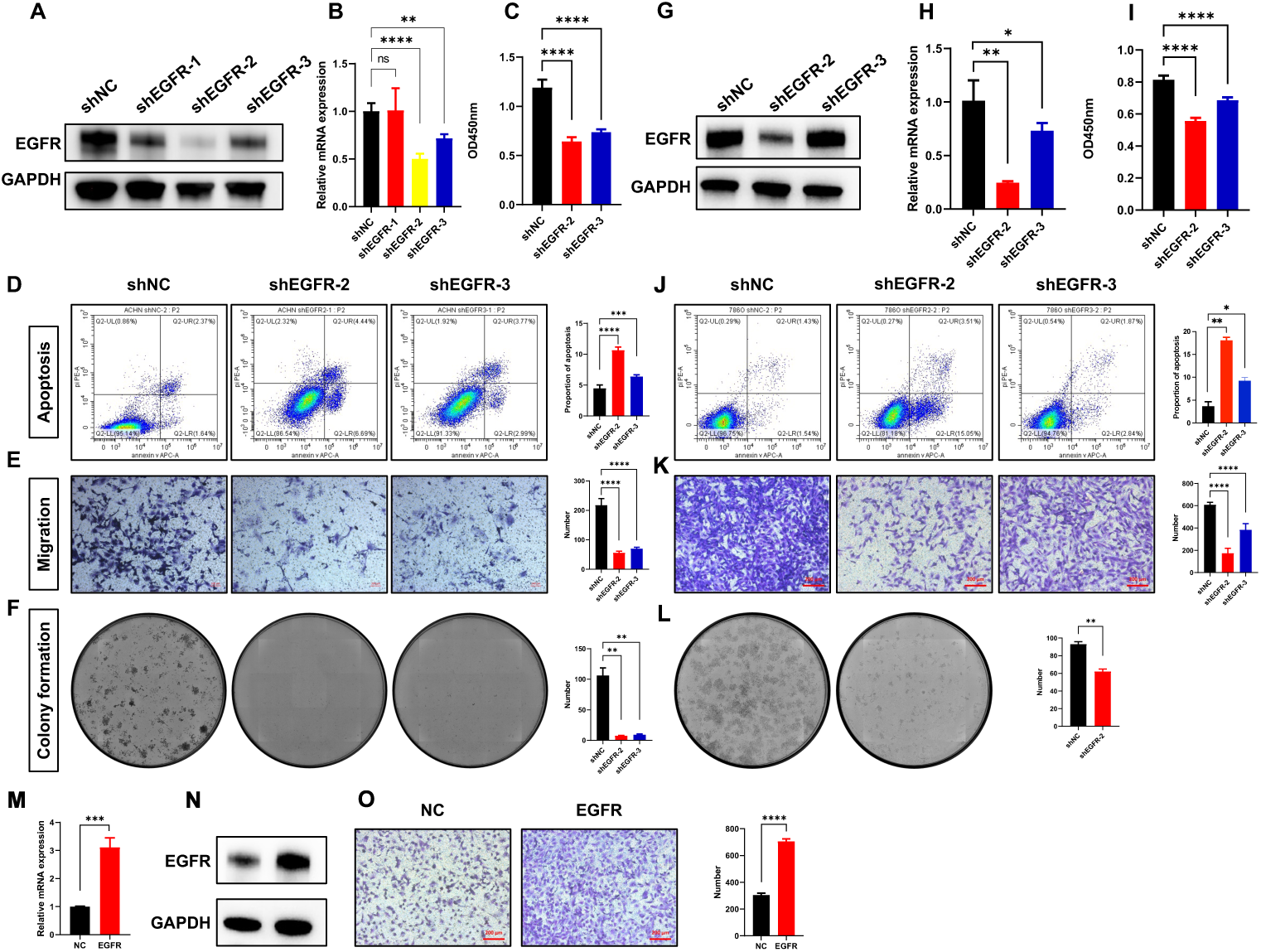
EGFR dysregulation significantly affect ccRCC cell progression *in vitro*. **A, B**. The protein and mRNA expression of EGFR in stable cells of ACHN-shEGFR. **C**. The cell viability of ACHN-shEGFR cells. **D, E, F**. The assays of apoptosis, migration, and colony formation in ACHN-shEGFR cells. **G, H**. The protein and mRNA expression of EGFR in stable cells of 786-O-shEGFR. **I**. The cell viability of 786-O-shEGFR cells. **J, K, L**. The assays of apoptosis, migration, and colony formation in 786-O-shEGFR cells. **M, N**. Overexpression of EGFR in 786-O cells at mRNA and protein levels. **O**. The migration assay of 786-O-EGFR cells. NC, negative control; ns, no significant; *, p-value*<*0.05; **, p-value*<*0.01; ***, p-value*<*0.001; ****, p-value*<*0.0001.

**Fig. 5.**
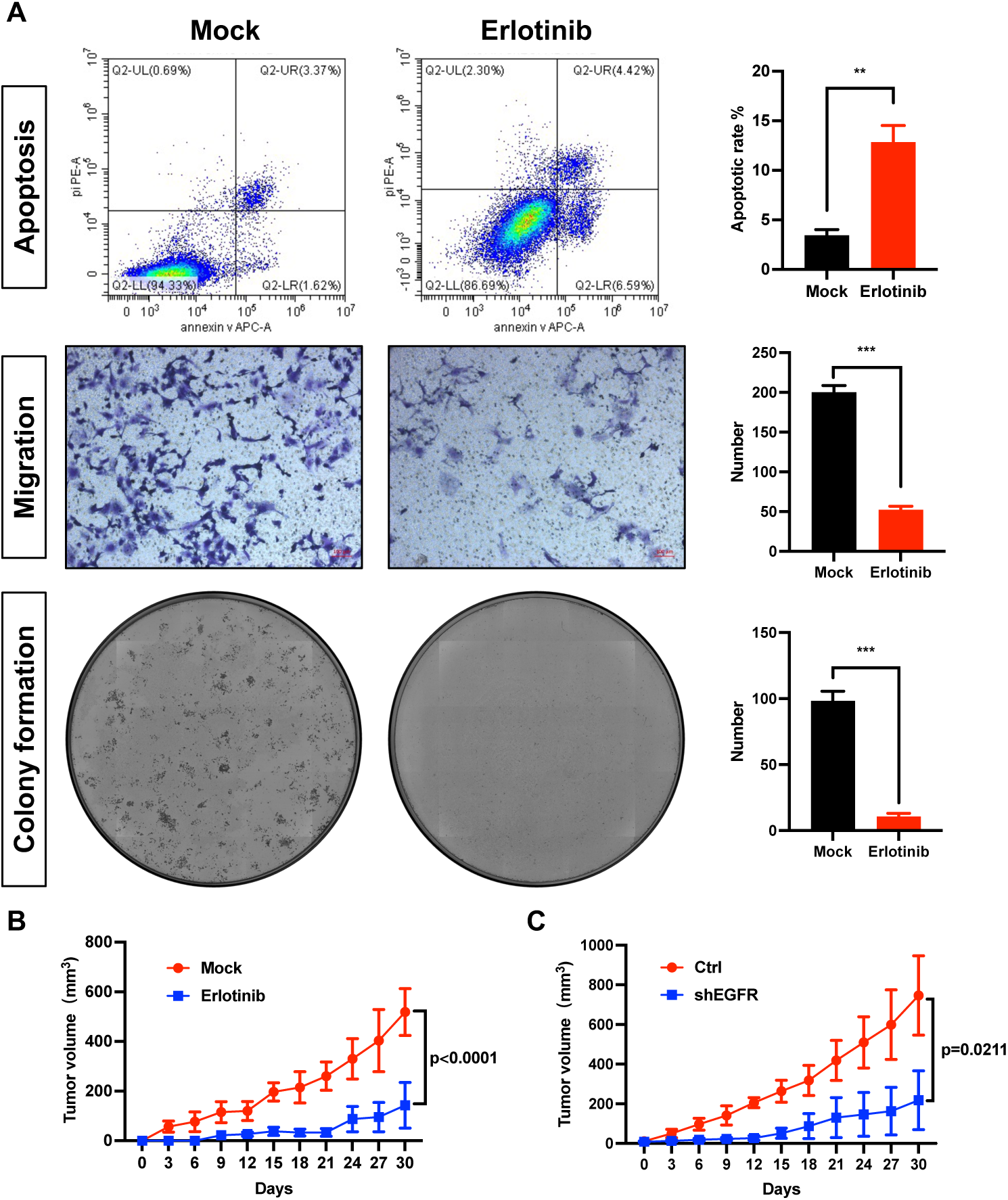
EGFR inhibition represses ccRCC cell progression *in vitro* and *in vivo*. **A**. Erlotinib markedly promoted apoptosis and inhibited migration and colony formation of ccRCC cells. **B, C**. Both erlotinib (B) and EGFR knockdown (C) significantly retarded tumor growth *in vivo*. **, p-value*<*0.01; ***, p-value*<*0.001.

## 3 Discussion

In this study, we developed RL-GenRisk, a novel approach utilizing deep rein-forcement learning to enhance ccRCC risk gene identification by integrating network knowledge with gene mutation data. By considering the risk gene identification as a node selection problem, we model the ccRCC risk gene identification as a Markov decision process, reducing the dependency on labeled data. Furthermore, we designed a data-driven reward and employed the DQN algorithm to optimize RL-GenRisk. RL-GenRisk exhibits a substantial improvement in the task of ccRCC risk gene identi-fication and reveals several novel risk genes. Moreover, since mutation data for patients with various types of cancer are now readily available, RL-GenRisk can be applied to other types of cancer in the future.

RL-GenRisk successfully identified known ccRCC risk genes. Among the top 20 ccRCC risk genes identified by RL-GenRisk, 10 genes are listed in the IntOGen database and recognized as ccRCC risk genes in prior research, including VHL [66, 67], PBRM1 [68, 69], SETD2 [70, 71], BAP1 [72, 73], MTOR [74, 75], ATM [76, 77], SPEN [78], and PTEN [79, 80]. This demonstrates that RL-GenRisk can effectively identify ccRCC risk genes. Interestingly, 10 of the top 20 identified ccRCC risk genes are novel and not listed in the IntOGen database. However, some recent studies have found that among these 10 novel genes, PDE4DIP, FLG, and SMARA4 are associated with ccRCC. For example, methylation levels of PDE4DIP were found to be associated with reduced overall survival in ccRCC patients [81]. FLG was found to be specifically mutated in specific subtypes of ccRCC [82]. SMARA4, a component of the SWI/SNF complex, was found to be frequently altered in ccRCC [83]. These latest findings validate the novel risk genes identified by RL-GenRisk. While some genes within these novel identified candidates have been shown a correlation with ccRCC, their molecular mechanism within ccRCC needs to be further investigated.

Furthermore, we validated a high-confidence ccRCC risk gene EGFR identified by RL-GenRisk with independent datasets and biological experiments. EGFR exhibits significant upregulation at both bulk and single-cell levels in ccRCC patients. The overexpression of the protein encoded by EGFR is significantly associated with poor survival of ccRCC patients. Moreover, a recent study [84] has found that treatment with the PLOD2 inhibitor minoxidil significantly inhibits the progression of ccRCC by inactivating the EGFR/AKT signaling axis. Furthermore, through comprehensive *in vitro* and *in vivo* experiments, we validated the impact of EGFR on ccRCC progression, demonstrating that decreased EGFR expression promotes ccRCC cell apoptosis and suppresses colony formation. Additionally, the use of the EGFR inhibitor erlotinib effectively inhibits ccRCC cell migration and growth in mice, highlighting the potential therapeutic significance of our findings.

## 4 Methods

### 4.1 Data preparation

Consistent with previous studies [29, 30], we used the information from the PPI network together with gene mutation data from patients. The feature matrix *H* for nodes is constructed based on the topological information of genes in the PPI network, the mutation frequency of genes in ccRCC patients, and the genes’ length.

#### PPI networks

We collected protein-protein interactions from HPRD [45], STRING-db [46], Multinet [47], IRefIndex [48], and HumanNet [49]. Then, following the previous study [30], we performed a two-step preprocessing on these PPI networks. First, we excluded the nine longest genes (TTN, MUC16, SYNE1, NEB, MUC19, CCDC168, FSIP2, OBSCN, GPR98) as they tend to acquire numerous mutations by chance and cover many patients [30]. Second, to mitigate the noise due to the dense connectivity in PPI networks, we applied the diffusion state distance (DSD) metric [85] on these PPI networks.

#### Gene mutation data

We collected somatic mutation of genes in ccRCC patients from The Cancer Genome Atlas, including 379 ccRCC patients and 14070 genes. Each gene corresponds to a patient list containing patients who carry mutations in that gene.

#### Feature representation

The initial feature of gene *v* is represented as 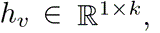 where *k* is the dimension. In RL-GenRisk, the node feature is flexible and can have different dimensions. We set *k* = 3 in RL-GenRisk. Therefore, the initial feature of gene *v* can be represented as 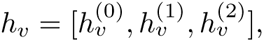 which respectively measures the topological importance of *v* in the PPI network, the mutation frequency in patients, and the length information of the gene. We first use the degree of *v* as the first dimension of *h_v_* since it can represent the topological importance of *v* in the PPI network:

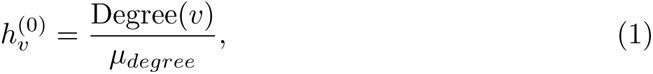

where Degree(*v*) represents the node degree of gene *v* in the PPI network. *µ_degree_* is a hyperparameter for normalization. We set *µ_degree_* equal to the maximum degree of nodes in the PPI network. Then, the second dimension of *h_v_* considers the mutation frequency of gene *v* in ccRCC patients:

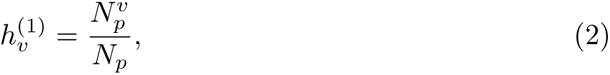

where 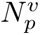 represents the number of patients carrying mutation on *v* and *N_p_* represents the total number of patients. Moreover, the initial feature should consider the length information of the gene, since the longer genes tend to include more mutations by change. To dilute this type of effect, the third dimension of *h_v_* is designed as follows:

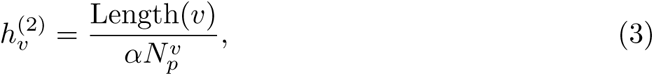

where Length(*v*) represents the gene length of *v*, and *α* is a normalization parameter. We set *α* equal to the node number of a connected subgraph in the PPI network that each patient carried mutations on at least one gene in this subgraph. Finally, the initial feature of gene was defined as 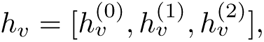 and the initial feature matrix for all genes was represented as 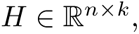 where *n* is the number of genes.

### 4.2 Key elements in RL-GenRisk

In RL-GenRisk, the identification of ccRCC risk genes is framed as a Markov decision process. At each step, the policy of RL-GenRisk receives the current state as input and selects an action. The action is selecting a node that connects with the sampled subgraph and appending this node to the sampled subgraph. Thus, this sampled subgraph adds a new node at each step. A reward is obtained after taking an action. Therefore, there are three key elements in RL-GenRisk, including state, action, and reward. These three key elements are defined as follows:

#### State

The state *s* at step *t* is represented as *s_t_*, which consists of the feature matrix *H* and the current subgraph *G_t_* sampled from the PPI network. In the first step, RL-GenRisk creates an empty subgraph and randomly adds a node to it. The design of incorporating the sampled subgraph into the state allows RL-GenRisk to delve into localized network structures. This enables RL-GenRisk to focus on the inter-action relationships relevant to the current sampled subgraph at each step. Therefore, the subgraph information in the state enables RL-GenRisk to accurately estimate its current environment and adjust its actions accordingly.

#### Action

At step *t*, the action *a_t_* is selecting a node that connects with the sampled subgraph *G_t_* and appending this node to *G_t_*. The action space at step *t* is represented as *A_t_*, which contains nodes that connect with *G_t_*. In detail, after getting *Q* values for all possible actions, RL-GenRisk uses an *ɛ*-greedy strategy to select an action *a_t_*:

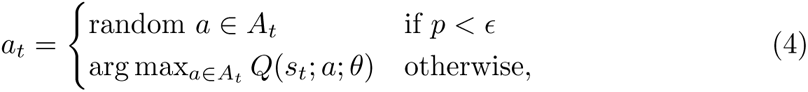

where *Q*(*s_t_*; *a*; *θ*) represents the *Q* value of action *a* calculated by the policy based on the current state *s_t_*, *θ* stands for the parameters of the policy. *A_t_* represents the action space at step *t*. With a probability *p* not exceeding *ɛ*, we select an action in action space randomly. Otherwise, the action with the highest *Q* value is selected. The *ɛ*-greedy strategy can enhance the exploratory nature of the policy, effectively preventing the model from getting stuck in local optimal during training. Same with the previous study [86], we set *ɛ* equal to 0.95. To balance exploration and exploitation, *ɛ* is decreased gradually during the training process.

#### Reward

In reinforcement learning algorithms, the design of the reward is crucial to the performance of the algorithm. In this study, we designed a specific data-driven reward based on the current sampled subgraph. The sampled subgraph *G_t_* at step *t* is expected to cover more patients so that risk genes are more likely to appear in it. Instead of focusing on an individual gene, RL-GenRisk focuses on a sampled sub-graph *G_t_* with node feature matrix *H*. This makes RL-GenRisk effectively leverage information from both the PPI network and gene mutation data. More importantly, this design of the reward ensures the accurate identification of genes with high mutation frequencies, and also identifies potential risk genes with low mutation frequencies but functionally interacting with genes that have high mutation frequencies. However, only considering the number of patients covered by the sampled subgraph *G_t_* is not enough. Longer genes are more likely to mutate by chance in patients. Therefore, shorter genes with high mutation frequencies are more likely to be risk genes [30]. To take gene length into account, we integrate gene length information when designing the reward. Thus, the single-step reward is higher when the sampled subgraph covers more patients and the genes in the subgraph have shorter lengths. The single-step reward *r_t_* at step *t* is designed as:

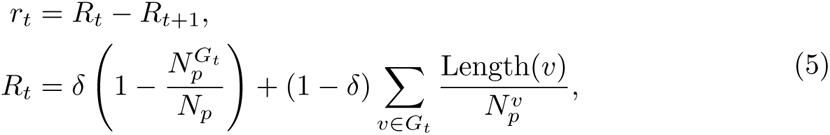

where *r_t_* represents the single-step reward at step *t*. *δ* is a weight hyperparameter. *R_t_* provides an evaluation score of the current state, with 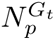 representing the number of patients that carry at least one mutation on the genes in the sampled subgraph 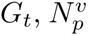 represents the number of patients carrying mutations on gene *v*, and *N_p_* represents the total number of patients. Length(*v*) represents the length of gene *v*. The cumulative reward is defined as the sum of all single-step rewards.

### 4.3 Policy network

In RL-GenRisk, the policy takes the current state as input and outputs *Q* val-ues for all possible actions. The policy of RL-GenRisk is represented by a neural network which is usually referenced as the policy network. Specifically, the policy of RL-GenRisk comprises two main components: a graph convolutional network (GCN) and a node evaluation network. We also use three multi-layer perceptrons (MLP) to perform dimensionality transformation. The GCN aggregates neighborhood informa-tion to get the representation for each node by multiplying the graph laplacian [41] with the node feature matrix. Given the feature matrix 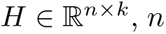 is the number of nodes and *k* is the dimension, the hidden representation of the *l*-th graph convolutional layer is calculated as:

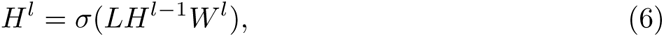

where *H^l^* represents the hidden representation of the *l*-th graph convolutional layer. 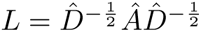 represents the normalized graph laplacian, which is used to aggregate neighborhood information. GCN preserves the original node signal by adding self-connections: *Â* = *Ã* + *I*, *Ã* represents the adjacency matrix and *I* represents the identity matrix. 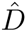 represents the degree matrix for *Â* and 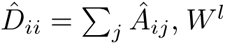 represents the trainable matrix of the *l*-th graph convolutional layer. *σ* represents the non-linear activation function.

In RL-GenRisk, we use two graph convolutional layers. Before the first graph convolutional layer receives data, we use an MLP to perform dimensionality transfor-mation on the initial features:

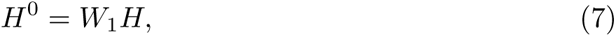

where *H* represents the initial feature matrix, *W*_1_ represents a trainable projection matrix. *H*^0^ represents the input feature matrix of the first graph convolutional layer. Inspired by residual networks [87], we concatenate the hidden representation matrices of two graph convolutional layers with *H*^0^, and use another MLP for dimension transformation to obtain the final representations matrix *Ĥ*:

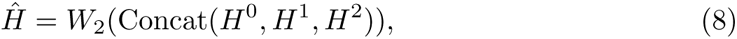

where *H*^1^, *H*^2^ represent the hidden representation matrices of the two graph convo-lutional layers, *W*_2_ represents a trainable projection matrix, and Concat(·) represents the concatenation operator.

At step *t*, after getting the representations matrix by GCN, RL-GenRisk cal-culates the representation of the sampled subgraph through the averaging pooling operation and uses the MLP for dimension transformation:

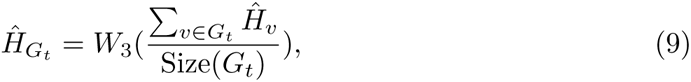

where 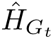 represents the representation of sampled subgraph *G_t_*, *Ĥ_v_* represents the representation of gene *v*, Size(·) represents the number of nodes in the sampled subgraph, and *W*_3_ represents a trainable projection matrix.

Then, we use a two-layer MLP as the node evaluation network to calculate the *Q* values for each action in action space:

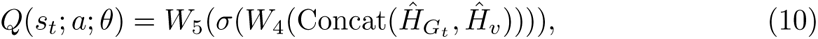

where *Q*(*s_t_*; *a*; *θ*) represents the *Q* value for action *a* based on the current state *s_t_*, *θ* stands for the parameters of the policy. 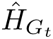 represents the representation of sampled subgraph *G_t_*, *v* represents the gene that selected by action *a*, and *Ĥ_v_* represents the representation of gene *v*. *W*_4_ and *W*_5_ represent two trainable projection matrices. *σ* represents the non-linear activation function. By using a two-layer MLP that can continuously update parameters during the training process as the node evaluation network, RL-GenRisk can better predict *Q* values for actions. More details about the policy network can be found in Supplementary Table 3.

### 4.4 Policy training for high-confidence risk gene identification

Consistent with the previous study [30], we sampled the training data, randomly collecting 85% samples from 379 ccRCC patients before the training process started. In the training process, since we model the ccRCC risk gene identification as a Markov decision process, the policy of RL-GenRisk iteratively receives the current state as input, selects an action, and obtains a reward. The DQN algorithm [42], which is widely used in reinforcement learning methods, is employed to train the policy of RL-GenRisk. DQN uses two sets of *Q* values calculated by an online network and a target network, respectively. The online network in our study is the neural network of RL-GenRisk. The target network is designed to prevent the online network from overestimating *Q* values. The architecture of the target network is the same as that of the online network, but their parameters are different. In RL-GenRisk, the loss function is defined as follows:

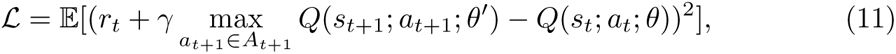

where *L* represents the loss to be minimized. *r_t_* stands for the reward received at the step *t*, and *γ* is the discount factor. 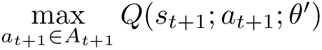 represents the target network estimate of the maximum expected *Q* value for the next state *s_t_*_+1_ and pos-sible action *a_t_*_+1_. *A_t_*_+1_ represents the action space. *Q*(*s_t_*; *a_t_*; *θ*) represents the *Q* value calculated by the online network based on the current state *s_t_* and action *a_t_*. *θ* and *θ^′^* represent the parameters of the online network and target network, respectively. *θ* is updated through gradient backward based on the loss function. Then *θ*^′^ is updated through soft updates based on the parameters of the online network as follows:

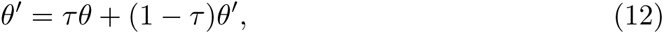

where *τ* represents a hyperparameter that controls the proportion of each update. More details about the hyperparameters in RL-GenRisk are shown in Supplementary Table 4.

In the identification process, RL-GenRisk differs from existing methods that predict outcomes following the same procedure as the training process [23, 29, 30]. Instead, capitalizing on the advantage of RL-GenRisk combined with reinforcement learning, we have designed a concise and effective identification process. Specifically, the model with the highest cumulative reward during the training process is selected as the best model and loaded. The sampled subgraph is initialized as empty, and all nodes in the PPI network are included in the action space at the beginning. The policy calculates *Q* values for all genes, and subsequently, the final ranking list of ccRCC risk genes is ordered based on these *Q* values. The higher *Q* value indicates higher risk. We analyzed the top 20 high-confidence risk genes in our study:

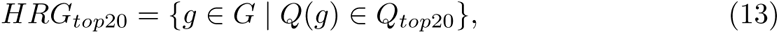

where *HRG_top_*_20_ represents the top 20 high-confidence risk genes. *g* represents a gene and *G* represents the PPI network. *Q*(*g*) represents the *Q* value of gene *g* and *Q_top_*_20_ represents the top 20 highest *Q* values in the set of all *Q* values calculated at the identification process. Researchers can select the number of top genes for verification based on actual verification costs.

### 4.5 Performance evaluation

To evaluate the results of all tested methods, we used a list of known ccRCC risk genes from the IntOGen database, including 27 ccRCC risk genes. The performance was evaluated using the Discounted Cumulative Gain (DCG), which has been used for evaluating cancer risk gene identification in the previous study [29]. The DCG is calculated as:

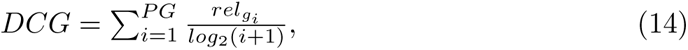

where *PG* represents the ranking list of the identified ccRCC risk genes. 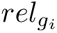 is equal to 1 if the *i*-th gene *g_i_* is contained in the IntOGen database, and 0 otherwise. Therefore, the higher the ranking of known ccRCC risk genes in the prediction results, the higher the DCG score. Following the previous study [30], the top 100 identified ccRCC risk genes were considered in performance evaluation. Additionally, to provide a more comprehensive evaluation of different methods, we presented DCG curves and calculated the area under the DCG curve.

### 4.6 Statistical analysis

#### Gene set enrichment analysis

We used the g:Profiler [54] for running functional enrichment analysis of the top 20 high-confidence risk genes identified by RL-GenRisk. g:Profiler maps genes to known functional information sources and detects statistically significantly enriched terms. We performed enrichment analysis on Gene Ontology, Human Phenotype Ontology, and WikiPathways. We used FDR p-value *<* 0.05 as the significance threshold.

#### Survival analysis

Clinical data of ccRCC patients and protein expression data of EGFR were obtained from TCGA. We used cSurvival [88] to perform progression-free survival and disease-specific survival analysis on these data. Progression-free survival utilizes the time from randomization or initiation of treatment to the occurrence of disease progression or death [64]. Disease-specific survival refers to deaths caused specifically by a particular disease [65]. Patients were categorized into quartiles based on the expression levels of protein encoded by EGFR. The Kaplan-Meier estimator was used to generate survival curves, and the difference was assessed using the log-rank test [89].

### 4.7 *In vitro* and *in vivo* experiments

#### Cell lines

ACHN and 786-O cells were purchased from Shanghai Zhong Qiao Xin Zhou Biotechnology Co., Ltd. (Shanghai, China), and were cultured in DMEM and RPMI-1640 medium (HyClone, Utah, USA), respectively, supplemented with 10% fetal bovine serum (FBS, Gibco, Australia) and 1% antibiotics (penicillin and streptomycin, HyClone) in a humidified incubator containing 5% CO2 at 37 °C. The stable cell lines of ACHN-shEGFR, 786-O-shEGFR, and 786-O-EGFR were obtained as described previously [90].

#### Western blot analysis

Total protein was extracted using TPER solution from Thermo Fisher Scientific. The protein concentration was measured using a BCA kit from Pierce. The proteins were separated using 10% SDS-PAGE. The proteins were transferred onto a PVDF membrane, and the membrane was blocked with 5% skimmed milk for 1 h at room temperature (RT). Then, the membrane was incubated with primary rabbit anti-human EGFR antibodies (R22778, ZenBio) at 4°C overnight. After three washes with TBST for 10 min, the membrane was incubated with horseradish peroxidase (HRP)-conjugated goat antirabbit secondary antibodies (458, MBL) for 1 h at RT. Finally, the protein bands were detected using an HRP substrate, and the intensity of the bands was quantified using ImageJ software. All experiments were performed in triplicate.

#### Quantitative polymerase chain reaction (qPCR)

Total RNA was extracted from the cells using a QIAGEN kit (74104), and the RNA concentration was determined using a NanoPhotometer from Implen (München). cDNA was synthesized using a RevertAid first-strand cDNA synthesis kit from Thermo Fisher Scientific (Waltham). PCR was performed using the following protocol: 98°C for 2 min and 40 cycles at 98°C for 5 s and 60°C for 10 s. The relative gene expression was calculated using 2*^−^*^ΔΔCt^. *β*-Actin was used as an internal control. All experiments were performed in triplicate.

#### Cell counting kit-8 assay

The cell counting kit-8 (CCK-8) assay was performed using a kit from Dojindo. A total of 3000 cells/well were cultured in a 96-well plate for 24 h, and 10 µL of CCK-8 solution was added to the wells. Then, the plate was incubated for 2 h at 37°C, and the optical density (OD) value was detected at 450 nm using a microplate reader. All experiments were performed in triplicate.

#### Cell apoptosi

An Annexin V-Alexa Fluor 647/PI Kit was purchased from 4A Biotech (Suzhou, China). The cells were digested and washed twice with cold phosphate-buffered saline (PBS). Next, the 1*×* binding buffer was used to suspend cells to a concentration of 1-5*×*10^6^ cells/mL. Then 5 µL of Annexin V/Alexa Fluor 647 was added to 100 µL of cells, and the mixture was incubated at RT for 5 min in the dark. Finally, the flow cytometry assay was performed after adding 10 µL of 20 µg/mL propidium iodide (PI) and 400 µL of PBS.

#### Transwell assay

Transwell migration assays were conducted using a Transwell chamber from Corning (REF 3422, Arizona, USA). Briefly, Transwell chambers were placed on a 24-well plate. Fresh medium containing 10% FBS in 600 µL was added to the lower chambers, and (2-5) *×*10^4^ cells in 200 µL of medium without FBS were added to the upper chamber. The 24-well plate was incubated at 37°C for 48 h. Cells that invaded through the chamber were washed, fixed (20 min with 4% paraformaldehyde), and stained (30 min with crystal violet). Then, the upper chambers were washed, photographed, and preserved under an inverted fluorescence OBSERVER D1/AX10 cam HRC microscope (Zeiss). Transferred cells were analyzed using ImageJ software.

#### Colony formation assay

The cells (500/well) were seeded into a 6-well plate and cultured at 37°C for 7-10 days. Then, the clones were imaged using a Celigo imaging cytometer from Nexcelom (Lawrence), and the clones were counted using ImageJ software. All experiments were performed in triplicate.

#### Animal study

BALB/C-nu/nu mice were purchased from GemPharmatech. A total of 5*×*10^6^ cells were inoculated into the right flank of mice, and the tumor volume was recorded every 3 days starting from day 17 after injection of the tumor cells. Mice were administered with erlotinib (S1023, Selleck, Shanghai, China) at a dose of 50 mg/kg. The tumor volume was calculated using the following equation: L×W^2^×0.5236, where L is tumor length and W is tumor width [90]. The animal procedures were approved by the ethics committee of West China Hospital, Sichuan University.

#### Statistical analysis

All data are presented as the mean ± standard error of the mean (SEM) or standard deviation (SD). Statistical significance for the comparison of multiple groups (*>*3) and between the groups was determined using analysis of variance (ANOVA) and Student’s paired t-test, respectively, in GraphPad Prism 9.0. p-value *<* 0.05 was considered statistically significant.

## 5 Data availability

All datasets analyzed in this article are publicly available. The HPRD network is available at http://www.hprd.org/. STRING-db is available at https://string-db.org/. Multinet and IRefIndex are available at https://github.com/raphael-group/hotnet2/tree/master/paper/data/networks. HumanNet is available at https://staging2.inetbio.org/humannetv3/. Mutations data of ccRCC patients are available at https://www.cbioportal.org/datasets. The clinical data of ccRCC patients from TCGA is available at https://xenabrowser.net/datapages/. The single-cell RNA-seq data of ccRCC patients is available at https://singlecell.broadinstitute.org/. Known ccRCC risk genes from the IntOGen database are available at https://www.intogen.org/search. Data for enrichment analysis is available at https://biit.cs.ut.ee/gprofiler/gost.

## 6 Code availability

The source code of RL-GenRisk and the trained model can be downloaded from the GitHub repository at https://github.com/23AIBox/RL-GenRisk.

## 7 Acknowledgements

This study was supported by the National Natural Science Foundation of China (92370132, 62072376, 82070784, and 62106172), the National Key R&D Program of China (2023YFC3403200), and the Xiaomi Young Talents Program of Xiaomi Foundation.

## 8 Author contributions

Conceptualization, J.P. and J.H.; methodology, J.P., J.H., Y.Z., D.L., Z.L., and L.H.; experimentation, D.L., Y.Z., Z.L., L.H., and J.A.; writing—original draft, D.L., and J.A.; writing—review and editing, J.P., J.H., J.A., D.L., Y.Z., X.Z., and S.J; supervision, J.P., J.H., and J.A.

