## Supplementary_Information for "Identifying potential risk genes for clear cell renal cell carcinoma with deep reinforcement learning"

### 1 Supplementary Figures

#### 1.1 DCG and AUC-DCG for different methods across different PPI networks

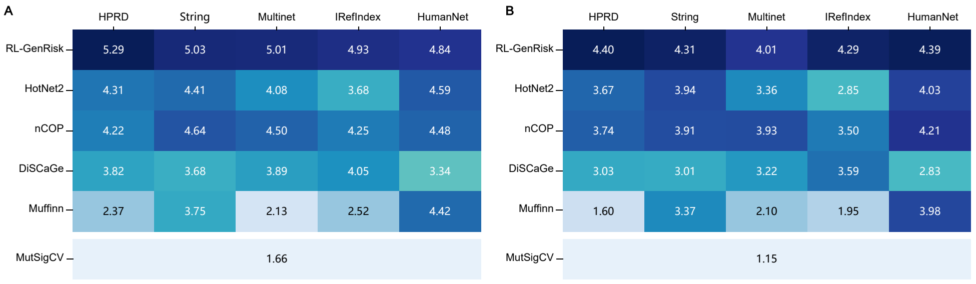

**Supplementary Fig 1**: **A**. Discounted Cumulative Gain (DCG) values for different methods across different PPI networks by top 100 predicted results. Dark blue cells in the heatmap correspond to high performance (high DCG scores), whereas light blue cells correspond to lower performance. **B**. Area under the DCG curve (DCG-AUC) for different methods across different PPI networks, divided by 100 for convenience.

#### 1.2 DCG curves for different methods across different PPI networks

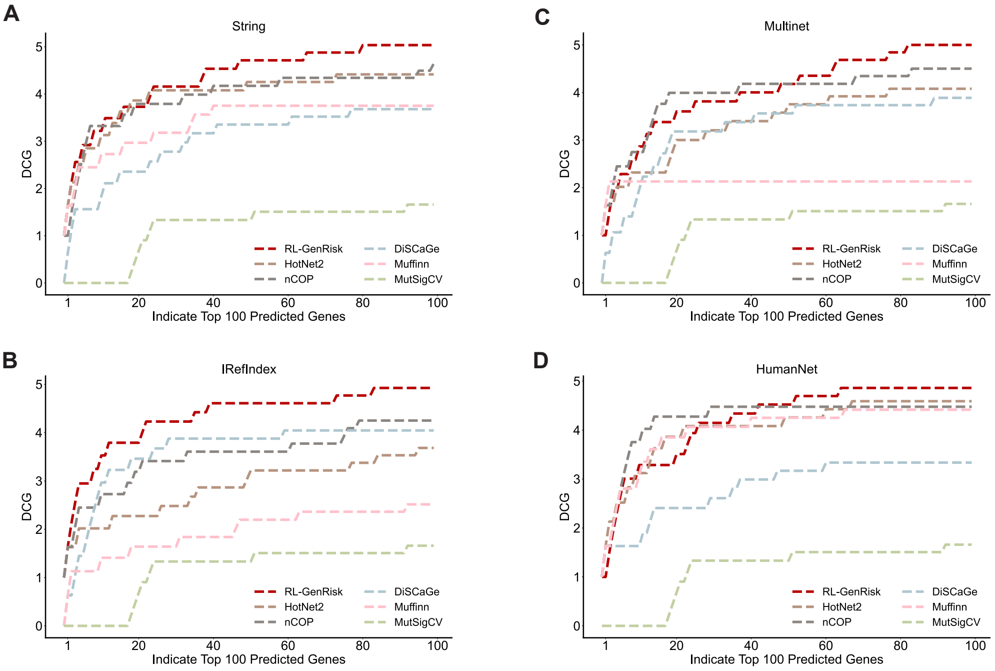

**Supplementary Fig 2**: Discounted cumulative gain (DCG) curves of the top 100 risk genes identified by different methods across different PPI networks, respectively.

#### 1.3 Mutation analysis of top 20 HRGs

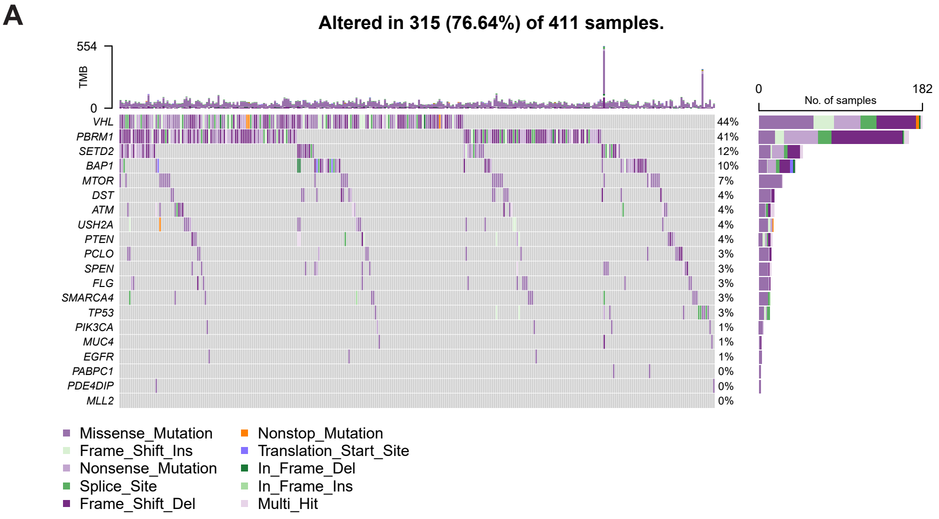

**Supplementary Fig 3**: **A**. Oncoplot of mutation data of ccRCC patients from TCGA obtained based on the top 20 high-confidence risk genes (HRGs). Each row represents a gene and each column represents a patient. Different colors represent different types of mutations.

#### 1.4 Enrichment analysis results on Human Phenotype Ontology

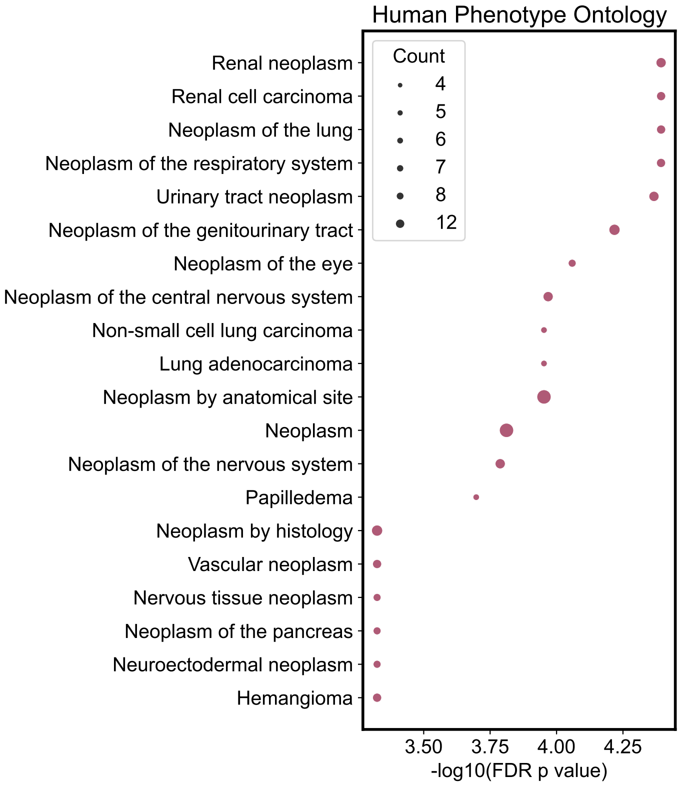

**Supplementary Fig 4**: Top 20 enriched items in Human Phenotype Ontology of top 20 high-confidence risk genes (HRGs) predicted by RL-GenRisk. The size of the circle corresponds to the number of top 20 HRGs contained in the corresponding item. The p values are adjusted by the False discovery rate (FDR).

#### 1.5 Expression levels of ten newly predicted genes in tumor and normal tissues

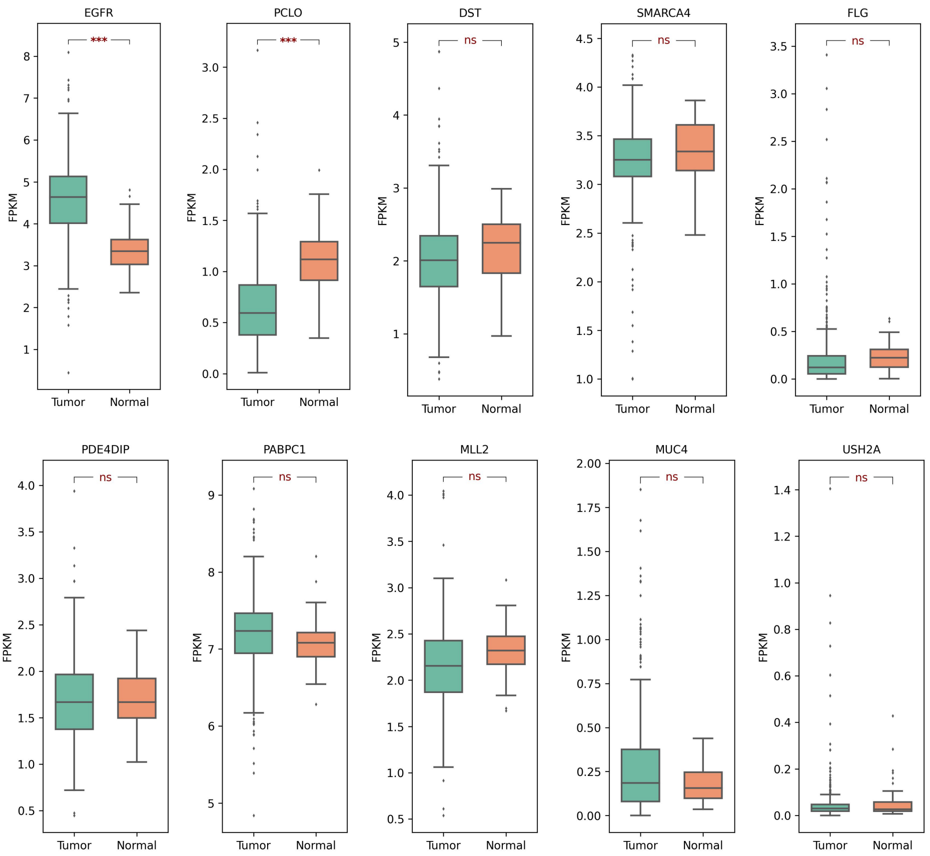

**Supplementary Fig 5**: Boxplots show the expression levels of the top 10 newly predicted HRGs (high-confidence risk genes) between tumor and normal tissues, using the RNA-seq data from 607 ccRCC patients in TCGA. The expression values are quantified using Fragments Per Kilobase Million (FPKM). ns, not significant; ***, p-value $< 0.001$

#### 1.6 Umap of expression valus of top20 HRGs in single-cell level of ccRCC patients

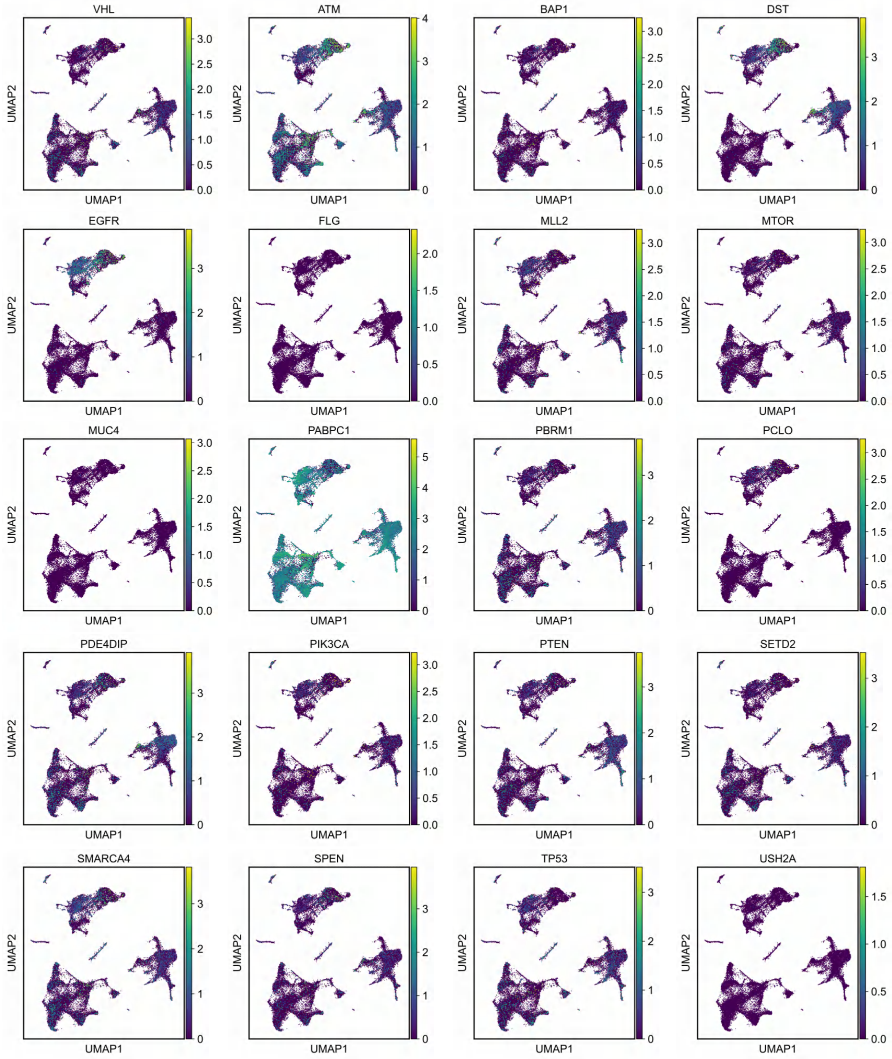

**Supplementary Fig 6**: Gene expression of top 20 high-confidence risk genes (HRGs) in different cells of seven ccRCC patients, with colors corresponding to the expression values.

#### 1.7 Violin plot of expression valus of top20 HRGs in single-cell level of ccRCC patients

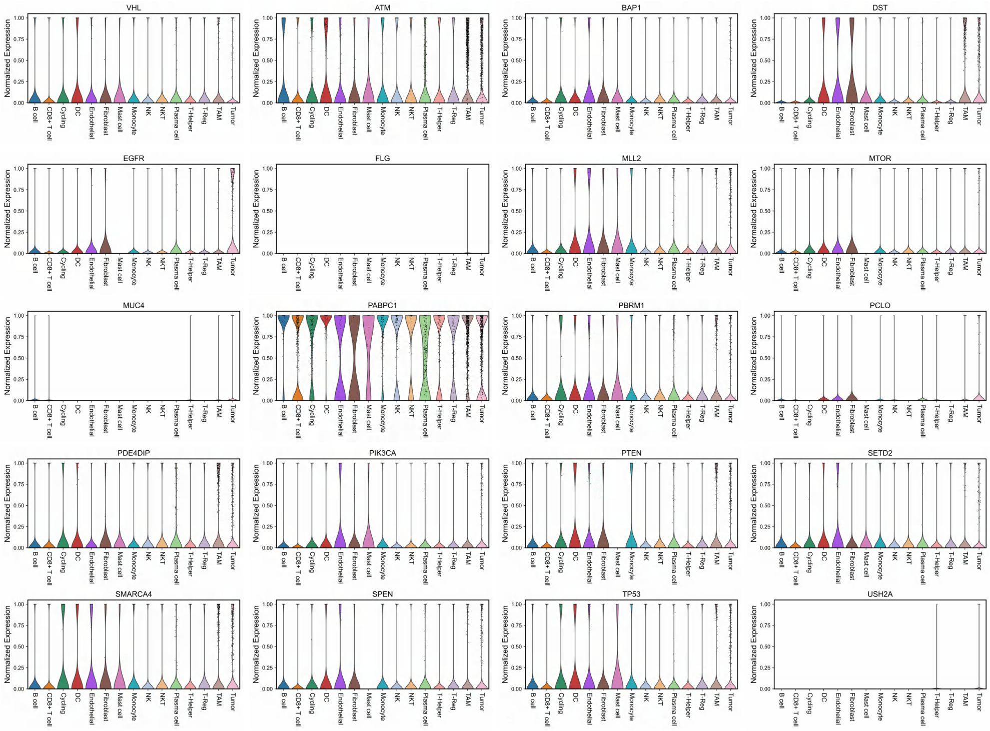

**Supplementary Fig 7**: Violin plots displaying the normalized gene expression of top 20 high-confidence risk genes (HRGs) in different cells of seven ccRCC patients in the single-cell RNA-seq data.

#### 1.8 Differential expression analysis at single-cell level

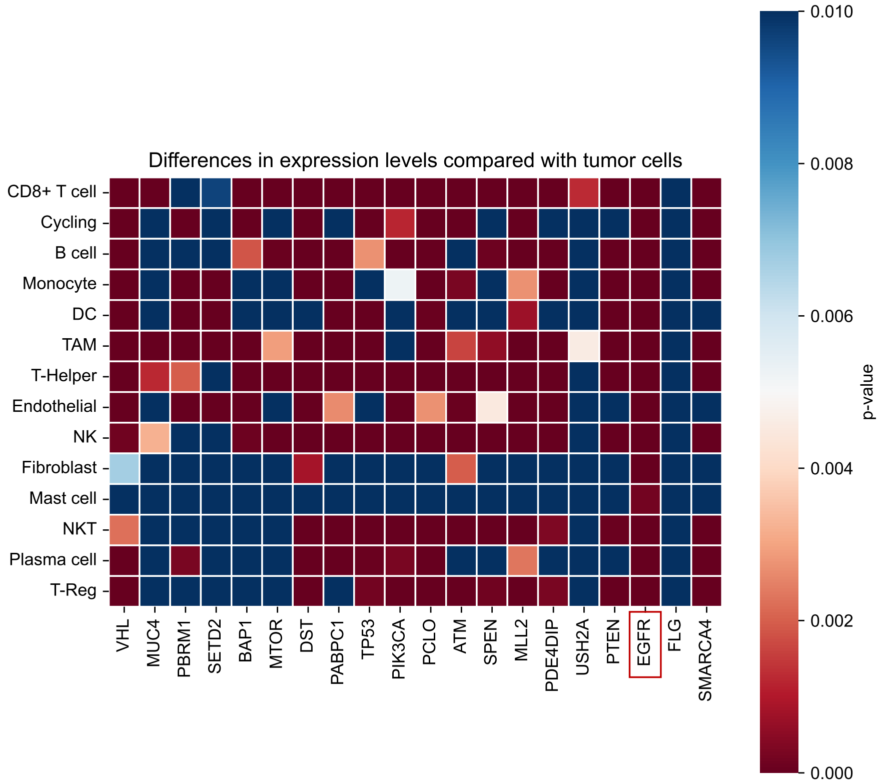

**Supplementary Fig 8**: Differential expression analysis at single-cell level. Each row represents a cell type, columns represent the top 20 high-confidence risk genes identified by RL-GenRisk. The color of the heatmap corresponds to the difference in expression of the gene in that cell type compared to tumor cells, using the Wilcoxon signed-rank test. The redder the color, the more significant the difference in expression at the single-cell level.

### 2 Supplementary Tables

#### 2.1 Top 20 HRGs identified genes by RL-GenRisk

**Supplementary Table 1**: The top 20 high-confidence risk genes (HRGs) identified by RL-GenRisk and whether genes are included in the IntOGen database.

| Ranking | Gene Symbol | Whether in IntOGen database |
|---|---|---|
| 1 | VHL | yes |
| 2 | MUC4 | no |
| 3 | PBRM1 | yes |
| 4 | SETD2 | yes |
| 5 | BAP1 | yes |
| 6 | MTOR | yes |
| 7 | DST | no |
| 8 | PABPC1 | no |
| 9 | TP53 | yes |
| 10 | PIK3CA | yes |
| 11 | PCLO | no |
| 12 | ATM | yes |
| 13 | SPEN | yes |
| 14 | MLL2 | no |
| 15 | PDE4DIP | no |
| 16 | USH2A | no |
| 17 | PTEN | yes |
| 18 | EGFR | no |
| 19 | FLG | no |
| 20 | SMARCA4 | no |

#### 2.2 Single-cell dataset information

**Supplementary Table 2**: Details of the single-cell dataset of ccRCC patients

| Dataset | Total number of cells | Number of genes | Cell types | Number of cells |
|---|---|---|---|---|
| Kidney from 7 ccRCC patients | 31856 | 32718 | Tumor cell | 7923 |
| | | | CD8+ T cell | 7668 |
| | | | B cell | 962 |
| | | | Monocyte cell | 1157 |
| | | | DC cell | 419 |
| | | | TAM cell | 5773 |
| | | | THelper cell | 3284 |
| | | | Endothelial cell | 271 |
| | | | NK cell | 2245 |
| | | | Fibroblast cell | 91 |
| | | | Mast cell | 39 |
| | | | NKT cell | 811 |
| | | | Plasma cell | 463 |
| | | | TReg cell | 750 |

#### 2.3 Structure of policy network in RL-GenRisk

**Supplementary Table 3**: Structure of policy network in RL-GenRisk

| Component | Layer detail | Input size | Output size |
|---|---|---|---|
| GCN | Linear | 3 | 64 |
| | GCNConv+ReLU | 64 | 64 |
| | GCNConv+ReLU | 64 | 64 |
| | Linear | 192 | 64 |
| Node evaluation network | Linear | 128 | 64 |
| | Linear+ReLU | 64 | 64 |
| | Linear+ReLU | 64 | 1 |

#### 2.4 Hyperparameters in RL-GenRisk

**Supplementary Table 4**: Hyperparameters in RL-GenRisk

| Hyperparameter | Default | Description |
|---|---|---|
| $\epsilon$ | 0.95 | the $\epsilon$ in the $\epsilon$-greedy algorithm |
| $\gamma$ | 0.95 | Reward decay rate |
| $lr$ | 0.001 | Learning rate for policy network during training using DQN algorithm |
| $\mu_{degree}$ | 180 | A weight for normalization for node feature |
| $\delta$ | 0.5 | A weight for calculating reward |
| $\tau$ | 0.001 | Control the synchronization speed of the parameters of the target network and the parameters of the policy network |
